# SUCCESSFUL *IN VITRO* EMBRYO PRODUCTION WITH OOCYTES ASPIRATED FROM LIVE WHITE-TAILED DEER (*Odocoileus virginianus texanus*) DONNORS UNDER CAPTIVITY IN NORTHEAST MÉXICO

**DOI:** 10.1101/2022.02.18.481106

**Authors:** Eduardo E. Maraboto M., Francisco J. Trejo M., Hilario Del Angel R., Juan Rosales H., Yuridia Bautista M., Libia I. Pérez T., Arnoldo González R.

## Abstract

Assisted reproductive technologies (ARTs), such as artificial insemination, semen sorting and freezing, embryo production *in vivo* and *in vitro*, are all methods in which animal reproduction management of several species has become considerably more efficient. These ARTs have been applied both in domestic and wild ruminants. In this study, *in vitro* embryo production was attempted with oocytes collected surgically during the mating season, from live white-tailed deer (WTD) hinds, maintained under captivity, in Northeast México. The study was conducted in two WTD farms nearby Cd. Victoria, Tamps., México, and another WTD farm located in Guerrero Coahuila, México. The laboratory work was carried at the Centro de Desarrollo de la Capacidad Productiva y Mejoramiento Genético de la Ganadería, property of the Unión Ganadera Regional de Tamaulipas (Centro-UGRT). The deer hinds are kept in captivity year-around, with feeding and health management provided accordingly to their requirements (50-60 kg females). Fresh green forage and clean fresh water is supplied daily. Oocytes were collected on each farm, while the hinds were maintained under general anesthesia and by means of a mid-ventral laparotomy, ovaries were exposed and follicles greater than 1-2 mm were aspirated with a 20 G needle, connected by a two-way plastic and latex hose to a vacum machine (WTA, Brazil); the oocytes were collected into a 50 ml. centrifuge tube containing wash media with heparine (Vitrogen, Brazil). Each oocyte collection lasted approximately 20 minutes, after which and while still on the farm, the oocytes were filtered with 50 µ mesh filters and rinsed several times with wash media, then placed on search Petri plates. Oocytes were then counted and classified (total, viable and non-viable oocytes) and placed into cryovials with maturation media (3 ml. MIV, Vitrogen, Brazil) and placed into a portable incubator with 5 % CO_2_ gas mix (LabiMixWTA, Brazil), after all hinds were done, oocytes were transported to the *in vitro* fertilization laboratory (IVF Lab) at the Centro-UGRT. Once at the IVF Lab, the oocytes were placed on a larger incubator (Eve, WTA, Brazil) and kept there for 18-20 hours, after which, the MIV media was changed for fertilization media (FIV, Vitrogen, Brazil) and fertilization was initiated by adding 10,000-12,000 live sperm per cryovial, and incubated for an additional period of 24 hours; after which, FIV media was replaced by the same media and at this point, cleavage rate was estimated by counting the oocytes that initiated cell división. At 72 hours after cell division started, fertilization rate was estimated; and 7 days after, the blastocysts were counted and classified. The whole process of oocyte maturation and embryo production *in vitro*, was conducted based on a beef cattle embryo production system and adapted to a deer embryo production system using media for small ruminants (Vitrogen, Brazil). Data collected per hind included total number of oocytes, viable and non-viable oocytes, cleavage rate (ratio of viable oocytes that initiated cell division over viable oocytes), fertilization rate (ratio of embryos that initiated cleavage over those that continued development to the blastocyst stage) and blastocyst rate (embryos reaching the blastocyst stage over cleaved embryos); averages were calculted for each parameter. The main results from this study on a per hind basis for total viable and non-viable oocytes were 9.8, 6.3 and 4.5, respectively; cleavage and blastocyst rates were 39.5 and 36.8, respectively and 2.3 blastocysts. In conclusión, oocyte collection from live WTD hinds and *in vitro* embryo production was succesfully done under farming conditions in Northeast México.

## INTRODUCTION

The deer family is a large group of ungulates, and the most evolved groups include the White-Tailed (WTD) and the Mule deer and the Black-Tailed deer, the WTD group includes more than 30 races or subspecies, of which more than one half arouse in México and other countries of Central and South America; thus, WTDs are presently found throughout the american continent (Taylor, 1969). During the last 4-5 decades, the WTD industry has developed into an important economic enterprise, where WTD bucks are the most sought trophy game by hunters (Green et al., 2017; Long et al., 2003; Saenz, 2007); in addition, both bucks and hinds are also sought for breeding purposes or other purposes (Berg and Asher, 2003; Long et al., 2003; Pintus and Ros-Santaella, 2014). Furthermore, expansion to farmed WTD enterprises have also evolved in support of the beef cattle industry, as a diversified enterprise (Cienfuegos et al., 2010).

Oocytes Collection from live donnor females by laparoscopic ovum pick up (LOPU) or surgical ovum pick up (SOPU, laparotomy), followed by *in vitro* fertilization (IVF) and *in vitro* embryo production (IVEP), as a single biotechnological tool has the potential of significantlly boosting the fast growing WTD farming enterprise in Northeast México (*Odocoileus virginianus texanus*), however, information is not readily available; which would allow a faster development of the industry, a fast genetic improvement, healthier status of herds and a broader biodiversity; by means of being a more effective method for *in vitro* embryo production, as it has been demonstrated in other small ruminants (Baldassarre et al., 1994; De Figueiredo and Magalhaes, 2010; Graff et al., 1999), beef cattle (Del Angel et al., 2020; González, 2020; Trejo et al., 2020) and in some other species of cervids (Comizzoli et al., 2001; García-Alvarez et al., 2011).

Earlier attempts for oocyte collection and IVEP in deer species have encountered some difficulties, mainly because of inconveniences such as availability, management and care of donnor hinds, since in most cases the donnors range freely in the wilderness; in consequence, oocyte collection for IVEP has been done postmortem in species as Red deer (Comizzoli et al., 2001; García-Alvarez et al., 2010), or by LOPU in Sika deer (Comizzoli et al., 2001), thus, assisted reproductive technologies (ARTs) must be adapted from other small ruminants (Rahman et al., 2008), including red deer (Berg and Asher, 2003); nevertheless, nowdays information on oocyte collection and IVEP in WTDs is not readily available, mainly due to difficulties in accessing biological materials and gametes, namely oocytes for IVEP. Thus, the objective of this study was to develop an IVEP method based on a beef cattle donnor IVEP system (Vitrogen, 2018) and using oocytes collected by SOPU from live WTD hind donnors kept in captivity and under the enviromental conditions in Northeast México.

## MATERIALS AND METHODS

The study was conducted at the IVF Laboratory of the Centro de Desarrollo de Capacidad Productiva y Mejoramiento Genético de la Ganadería Dr. Jorge R. Arnáez Gómez, property of the Unión Ganadera Regional de Tamaulipas (Centro-UGRT), located at the coordenates 23°44’06″ of latitude N and 99°07’51″ of longitude W and at 327 masl, the climate at the Centro-UGRT is BS1 (h’)hw, semi-dry with summer rains and a mean monthly rainfall of 62.3 mm, with mean maximum and mínimum summer temperatures of 45 and 20 º C, respectively and a mean relative humidity of 70% (INEGI, 2017).

The oocytes used for the study were collected from 7 (two hinds in Farm 1 and 5 in Farm 2) WTD adult hinds (two to three years old), property of two private WTD farms located within 10 miles of the Centro-UGRT and a third farm located 390 miles northwest of the Centro-UGRT; the hinds from Farms 1 &2 were kept under captivity and fed daily with fresh green buffel grass (*Cenchrus ciliaris spp*) or green sorghum forage *ad lib*., 2 kg of a comercial concentrate (containing 22% of crude protein) and a comercial mineral mix *ad lib*., fresh, clean water was also provided daily. The hinds were managed and kept in pens (30 m by 50 m) with wire mesh fence (3 m high) and were managed and cared according to the regulations of the Ethical and Animal Care Committee of the Facultad de Medicina Veterinaria y Zootecnia (Universidad Autónoma de Tamaulipas). The oocytes were collected during the months of december, 2021 and January 2022.

The oocyte collection was performed from each deer hind under general anesthesia, by a SOPU methodology, adapted from sheep, exposing the ovaries (mid-ventral laparotomy, González et al., 1987; Hulet and Foote, 1969; Rahman et al., 2008), a 3 cm incision was done in the mid-ventral line, immediately anterior to the udder, the ovaries were exposed individually and oocytes were collected using a vacum pressure machine (WTA, Brazil) and a plastic tubing fitted with a 2” long 20 G needle and connected to a 50 ml centrifuge tube, containing wash media (Vitrogen, Brazil), vacuum (50 to 80 mm Hg) was applied through a latex tubing connecting the vacuum machine with the centrifuge tube. Hind donnors were darted with the anesthetics (a mixture of 10 % Xilazine, Aranda Laboratories, México, plus Zoletil 100, Virbac, México), after the collection was done, the anesthesia was reversed (Tolazoline hidrochloride, Nexgen, U.S.A.). One centrifuge tube was used for each deer hind, the whole SOPU procedure lasted from 18 to 25 minutes, after which, the incision was closed, stiching separately the fascia and the skin, using # 1 G absorbable suture.

After each oocyte collection, the obtained oocytes were classified (total, viable and non-viable), the IVF procedure was adapted from the IVF beef protocol (Del Angel et al., 2020; Trejo et al., 2020), using all media for small ruminants (Vitrogen, Brazil); the oocytes from each collection tube were sieved with wash media, and filtered using 50 µ mesh filters, then the oocytes were transfered to search Petri dishes (Vitrogen, Brazil), where oocytes were located, and then transferred to incubating plates containing maturation media (Vitrogen, Brazil), the plates were covered with mineral oil (MIV, Vitrogen, Brazil) and placed into the incubator (Eve model, WTA, Brazil), at 38.5ºC, with a 5 % gas mix of CO_2_, for 24 hours, prior to fertilization. Initial oocyte search and classification (Vitrogen, 2018; Watanabe et al., 20017) was done at each farm, oocytes from each hind were placed into cryovials (3 ml, Vitrogen, Brazil) and incubated in a portable incubator (WTA, Brazil) and thus transported to the IVF laboratory, where they were placed in the Eve incubator (WTA, Brazil).

After maturation, oocytes and semen were prepared for fertilization, frozen (purchased or frozen at the Centro-UGRT) semen was prepared using a double-centrifugation method, after thawing, semen was placed in eppendorf centrifuge tubes containing Percoll (0.5 ml, Vitrogen, Brazil), the tubes were centrifuged during 12 minutes at 2500 G; after which, the semen pellet was placed in a second eppendorf tube, containing 0.25 ml of Percoll and 0.25 ml of fertilization media (Sperm, Vitrogen, Brazil) and centrifuged during 12 minutes at 2500 G. Prior to fertilization, oocytes were transfered to a new plate (one plate per donnor hind) containing fertilization media (FIV, Vitrogen, Brazil), and fertilization was done by adding 10-12000 live sperm (7 to 10 µl) per plate (initial sperm motility ranged from 60 to 70%); after addition of the sperm cells, plates were then placed into the incubator for a further period of 18-20 hours.

After the 18-20 hour fertilization incubation period, all oocytes were placed on new plates with embryo culture media (CIV, Vitrogen, Brazil), replacing the FIV media, and plates again were placed into the incubator for another 48 hours, at which time, cleavage rate was determined; at this time, the CIV media was replaced again. Furthermore, the embryos were incubated for a period of 96 hours. At the end of the final incubation period, embryo production rate and quality was estimated, using a stereoscope and the International Embryo Transfer Society standards (Wright, 1998); embryos were then vitrified (Del Angel et al., 2020; Trejo et al., 2020) for future transfer. Due to the low number of observations in the study, only basic statistics (means and percentages) data is presented.

## RESULTS AND DISCUSSION

The success of IVEP systems is largely dependent on the continuous supply of good quality oocytes, preferably from genetically valuable females, further, oocyte collection is also highly dependent on the species, in small ruminants, such as sheep (Baldassarre et al., 1994) or goats (Graff et al., 1999; Rahmann et al., 2008) and in some cervids, as Red (Berg et al., 1995; Locatelli et al., 2005) and Sika deer (Comizzoli et al., 2001; Locatelli et al., 2006) the most reliable method is by LOPU; although, other methods have been used successfully (see review by Rahmann et al., 2008.

In this study, oocyte collection was done by laparotomy (González et al., 1987; Hulet and Foote, 1969), and mean numbers for total and viable oocytes and number of blastocysts in this study were 9.8, 6.3 and 2.3, respectively; where as cleavage and blastocyst rate were 39.5% and 36.8 %, respectively. The results found here on all three farms are presented in Table 1. Based on the information available in several search platforms, this is the first report, where oocytes collected from live WTD hind donnors were used to produce embryos *in vitro*, during the mating season and under natural estrous cycle conditions and without hormonal estimulation.

**Table 1.**
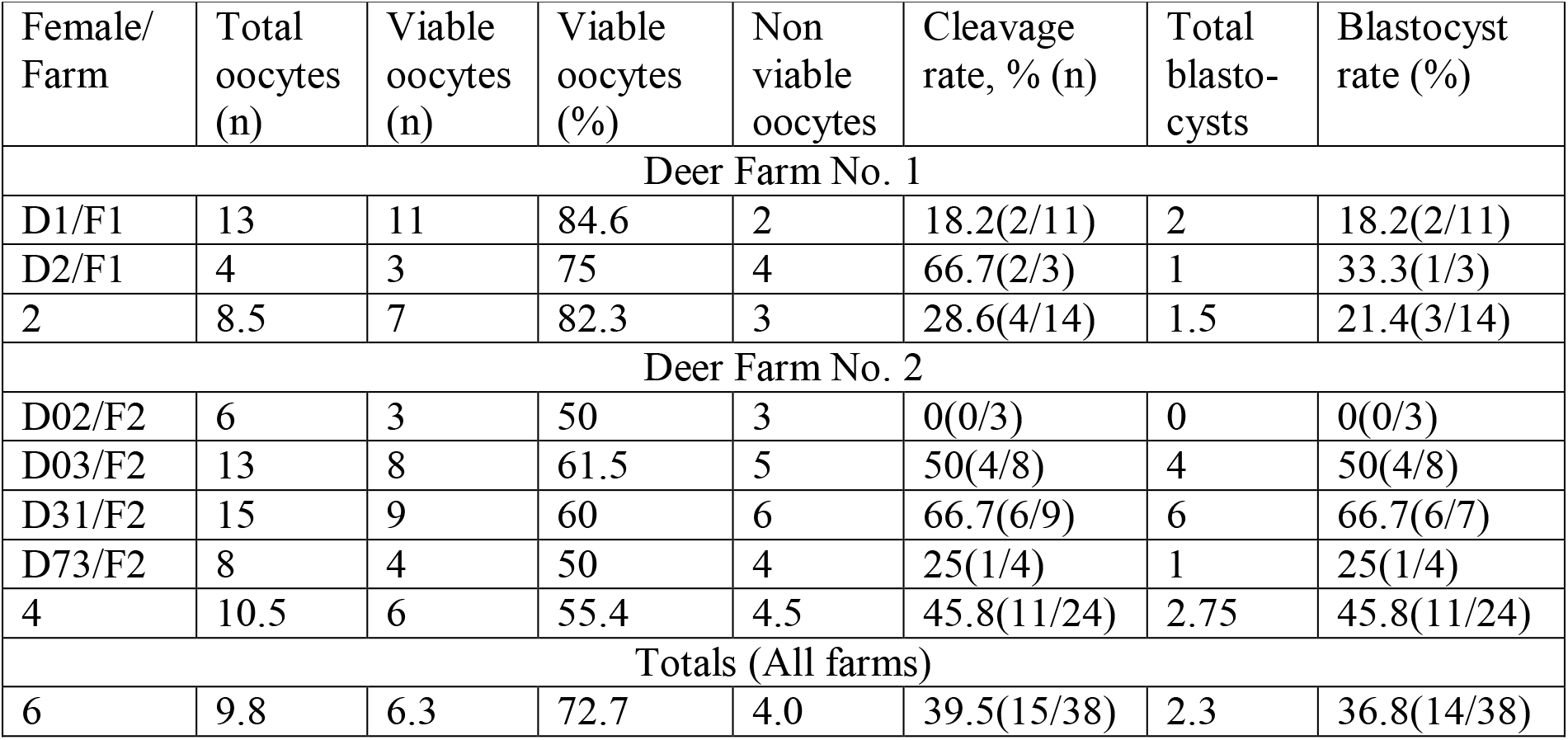
Individual and mean statistics for oocytes aspirated from White-Tailed deer hinds and blastocysts produced (Mean numbers and percent) *in vitro* in two deer farms under dry tropic conditions of Northeast México.

A wide variation in numbers and percentages of data was detected, both, in terms of farms used for the study and among deer hinds variability. However, mean numbers of total and viable oocytes and blastocyst rate were somehow similar in the three farms, these values were 8.5, 7 and 1.5 and 10.5, 6 and 2.75, respectively, for farm 1 and farm 2. On the other hand, the mean percentages for viable oocytes, cleavage rate and blastocyst rate were 82.3, 28.6 and 21.4 and 55.4, 45.8 and 45.8, respectively for farm 1 and farm 2 (Table 1). Noticeable differences were encountered in the number of blastocysts between farms, 1.5 (farm 1) *vs* 2.75 (farm 2); in addition, cleavage and blastocyst rates (Table 1) were greater in farm 2 (45.8 and 45.8 %) *vs* farm 1 (28.6 and 21.4 %).

Again, viable oocytes, blastocysts and blastocyst rate in this study were 6.3, 2.3 and 36.5 %, on a comparative basis, such results indicate that the oocyte and IVEP technology from cattle can be used to produce embryos from WTD cycling hinds; further, this is the first report for Northeast México and Southern United States, where collection of oocytes from live WTD hinds were used to produce *in vitro* embryos, with success rates similar to beef cattle.

On a comparative basis, the same numbers for beef cows are 20.1, 6.1 and 30.3 % (Trejo et al., 2020); from a technical point of view, the blastocyst rate was similar and in agreement with values reported in the literature for cattle managed under similar conditions (González, 2020; Watanabe et al., 2017). Results for other species, namely, sheep, goats and other cervids and from other studies following similar methodology for oocyte collection and IVEP will be discussed and compared with results from the present report.

There are two key issues regarding oocyte collection and IVEP for cervids as the WTDs, the first one is to have a constant, consistent and reliable source of good quality oocytes and in sufficcient quantities to make the IVEP process more productive; the second issue is to also have a consistent supply of good quality semen (fresh or frozen), factor which is no longer a problem, since freezing WTD buck semen has been done with good results (17 straws and 30 straws successfully obtained for this study) since some time ago (see review by Saenz, 2007). Semen from WTD bucks has been successfully frozen for IVEP in the semen laboratory of the Centro-UGRT.

Regarding a constant and consistent oocyte supply from WTD hinds, the available information is scarce and efforts to collect oocytes from small ruminants have been made using alternative collection methodologies; methods which include often times, post-mortem collection from hunting and abbatoirs, laparotomy (SOPU), LOPU or transvaginal OPU (TAGU, Berg and Asher, 2003; Graff et al., 1999; Rahman et al., 2008), recently, a video-LOPU system has been used at the Centro-UGRT for WTD hinds, which has been adapted from a sheep system (Teixeira et al., 2011). Besides, earlier reports indicated that cattle and sheep media and IVEP methodology were not suitable for deer IVEP (Berg and Asher, 2003). Further, in a recent report, the possibilities of applying ART technologies, such as gonadotropin therapy for ovarian stimulation for superovulation in wild animals was considered as non-viable in reference to increase the number of oocytes per sesión of OPU (Korzekwa and Kotlarczyk, 2021); although, a previous report documented the successful superovulation and embryo production from WTD hinds, where 13 transferable embryos were collected from six females (Waldhalm et al., 1989).

As shown in Table 1, blastocysts were successfully produced *in vitro* for the first time from oocytes collected from live WTD hinds, 5 out of 6 deer hinds produced embryos (with a range of 1 to 6 blastocysts per hind), the mean number of embryos per donnor hind was 2.3; in addition, both cleavage and blastocyst rates were similar for the two farms (29 and 46 % and 21 and 46 %, respectively), totals for both farms were 39 and 37 %, ranges for cleavage and blastocyst rates were 18 to 67 %, for both parameters and both farms. Similar information for WTD is not readily available, moreover, in an earlier publication, 2.3 embryos were reported to have been produced *in vivo* (Waldhalm et al., 1989). Blastocyst rate for goats and sheep have been reported to be of 50 and 61.5 % (Cox and Alfaro, 2007).

Information regarding oocyte collection and IVEP from several reports for different types of deer and other small ruminants, as goats and sheep, indicate similar numbers to those found in this study, both, in terms of viable oocytes and blastocysts. Teixeira et al. (2011) reported 6.3 oocytes collected from Santa Ines ewes, no embryo data was reported in this study. Data for goats has been reported to vary from 4.3 to 18 oocytes (Cognie, 1999, Graff et al., 1999, Samaké et al., 2000). Recent reports on LOPU for several species (Baldassarre, 2021, Neto et al., 2018) have indicated the advantages of collecting good quality oocytes for IVEP; oocyte crops vary from 10 to 15 oocytes (Baldassarre, 2021), whereas, the oocytes collected vary from 1.2 to 6 for deer and unpublished results for oocyte collection from WTD hinds of 19.8 oocytes (see review by Neto et al., 2018). Further, data for IVEP in these two latter reports was not provided. Earlier, LOPU was performed in Red deer, 44 oocytes were aspirated in 39 sessions, where 17 of 27 oocytes cleaved, embryos only developed until the 16 cell stage and no blastocysts were produced (Bainbridge et al., 1999). In another report with Sika deer, 3.9 oocytes per hind were collected by LOPU and blastocysts were produced *in vitro*, however, blastocyst data was not reported on a per hind basis (Locatelli et al., 2006); in this study, 2.3 blastocysts were produced, with a range of 1 to 6 blastocysts per hind. It should be pointed out that LOPU is usually conducted under ovarian stimulation, in this study, oocyte aspiration was conducted without ovarian stimulation.

## CONCLUSIONS AND FINAL IMPLICATIONS

Biotechnology includes several tools and its application has an enormous impact on domestic and wildlife production and management, ART’s as oocyte collection and IVEP are worth mention in more detail. In particular, IVEP has been applied with enormous benefits in cattle, goats and sheep; on the other hand, in wild species, as deer and other cervids, ART’s and in particular IVEP has been applied with unpredictable responses and varying results. The application of LOPU and IVEP has successfully been implemented in Red and Sika deer, in this study, OPU was performed by mid-ventral laparotomy from live WTD hinds, further, oocytes were matured *in vitro* and blastocysts were produced in vitro; oocyte and blastocyst numbers are modest, but promising and improvements should be made, particularly in the number of oocytes collected must be incremented, perhaps stimulating follicle growth with gonadotropins.

In wildlife and in particular for WTD’s production and management, LOPU and IVEP may be implemented under two scenarios, in the first one, under free wildlife conditions, as such, the results with ART’s will not match the results of the species under natural conditions because of its high gestation and fawning rates; In the second scenario, under intensive farming conditions and under captivity, the application of ART’s as LOPU and IVEP will allow improvement on the reproduction rates of genetically superior breeding does and bucks, which in turn, these does and bucks can be used of the same farm or they may be supplied to other farms or other production centers. The benefits of ART’s application in WTD’s will allow improve current production rates, will improve the economy of the regions where WTD’s are raised and will broaden biodiversity.

## ACKNOWLEDGEMENTS

The authors acknowledge the support provided by the two deer farmers that provided the deer hinds used for oocyte collection and the farm staff at each deer farm, for asistance provided. Acknowledgement is also expressed to all personnel at the Centro-UGRT that provided support and assistance for semen collection and freezing, the oocyte collection and the laboratory work for IVM and IVF conducted in this study, and acknowledge support provided by the Facultad de Medicina Veterinaria y Zootecnia of the Universidad Autónoma de Tamaulipas to conduct the present study.

